# Group optimization methods for dose planning in tES

**DOI:** 10.1101/2025.03.18.643934

**Authors:** R. Salvador, J. Zhou, B. Manor, G. Ruffini

## Abstract

**Objective:** Optimizing transcranial electrical stimulation (tES) parameters—including stimulator settings and electrode placements—using magnetic resonance imaging-derived head models is essential for achieving precise electric field (E-field) distributions, enhancing therapeutic efficacy, and reducing inter-individual variability. However, the dependence on individually personalized MRI-based models limits their scalability in some clinical and research contexts. To overcome this limitation, we propose a novel group-level optimization framework employing multiple representative head models.

**Approach:** The proposed optimization approach utilizes computational modeling based on multiple representative head models selected to minimize group-level error compared to baseline (no stimulation). This method effectively balances focal stimulation intensity within targeted brain regions while minimizing off-target effects. We evaluated our method through computational modeling and leave-one-out cross-validation using data from 54 subjects and analyzed the effectiveness, generalizability, and predictive utility of anatomical characteristics.

**Main results:** Our approach demonstrated that group optimization significantly outperformed protocols derived from standard templates or randomly selected individual models, notably reducing variability in outcomes across participants. Additionally, correlations between anatomical features (e.g., head perimeter and tissue volumes) and E-field parameters revealed predictive relationships. This insight enables further optimization improvements through the strategic selection of representative head models that are electro-anatomically similar to the target subjects.

**Significance:** The proposed group optimization framework provides a scalable and robust alternative to personalized approaches, substantially enhancing the feasibility and accessibility of model-driven tES protocols in diverse clinical and research environments.

**Data Access Statement:** The data that support the findings of this study are available from the corresponding author, R.S., upon reasonable request.

## Introduction

The primary biophysical mechanism through which transcranial electrical stimulation (tES), including transcranial direct current stimulation (tDCS), interacts with neural tissues is the induced electric field (E-field), which drives the observed neuromodulatory effects. Thus, effective tES intervention critically depends on optimal dose planning, encompassing all controllable stimulator parameters (current amplitude and waveform), electrode positioning and geometry, and stimulation timing (Peterchev et al., 2012). Early studies have highlighted the importance of carefully selecting these parameters to maximize therapeutic efficacy and minimize unwanted variability in stimulation outcomes (Miranda et al., 2013; Nitsche & Paulus, 2000; Saturnino et al., 2015). With the rapid evolution in the field of image processing and the advent of powerful computational resources, several tools have become available to create volume conductor models that represent the passive electrical characteristics of different head tissues from structural images, usually T1w-MRIs (Datta et al., 2009; Miranda et al., 2013; Windhoff et al., 2013). These biophysical head models were leveraged to optimize dose parameters such as electrode location and current, using optimization algorithms (Dmochowski et al., 2011; Fernández-Corazza et al., 2020; Ruffini et al., 2014; Saturnino, Siebner, et al., 2019). Despite differences in the optimization function that these algorithms minimize (or maximize), and the methods that are used to optimize said function, all algorithms share common steps: they rely on pre-computed solutions of the E-field distribution (lead-field) induced using predetermined electrode positions; they assume some model for the interaction of the E-field with the neurons that determines the choice of optimization function; and they impose constraints on the currents for safety considerations and hardware specifications. Initial studies using these approaches typically relied on non-personalized head models based on template MRIs (Splittgerber et al., 2020; Zhou et al., 2021). With advances in the processing pipelines, the use of head models personalized using subject-specific MRIs has become possible. This approach may improve stimulation effectiveness and lower its inter-subject variability (Salvador et al., 2021). Studies employing personalized, optimized protocols based on these algorithms have shown promising results in several recent trials (Daoud et al., 2022; Kaye et al., 2021; Splittgerber et al., 2021; Sprugnoli et al., 2019). However, the practical implementation of personalized head modeling remains challenging in several scenarios. For instance, in clinical trials, MRIs may be unsuitable for creating accurate head models because of technical limitations, such as inappropriate acquisition sequences, severe motion-related artifacts, or excessive crops of the scalp, skull, and/or cerebrospinal fluid (CSF) tissues. In addition, obtaining new images can be constrained by timing or budgetary constraints. Furthermore, structural T1 image processing may fail in certain populations, including those with lesions, genetic malformations, and other structural anomalies. Personalized head models are also incompatible with studies that require the montage and protocol to be predefined before the participant imaging data becomes available.

Here, we propose a group optimization algorithm that improves upon single non-personalized head model optimizations in terms of the optimization metric, specifically the proximity between the induced normal component of the electric field (En-field) and the target En-field distribution. In this approach, instead of using a single subject/template for optimization, a representative sample of multiple subjects from the target population is used in the optimization, which should, on average, reduce the penalty in the objective function of the optimization resulting from a standard template model. This approach has been used in studies on depression with positive results (Ruffini et al., 2024) and is currently being employed in two ongoing trials (see details below).

## Methods

### Comparison plan

To study the performance of group optimization we compare normalized error with respect to no intervention (NERNI) goodness-of-fit of group-optimized montages in our cohort with personalized and *standard template*derived solutions. Additionally, we compared these approaches against an approach in which the protocol was derived from one of the subjects in the cohort (non-personalized solutions). Standard templates are commonly used to determine non-personalized protocols in many studies, as are head models obtained from subjects with similar demographic characteristics to the population of interest.

### Head model creation

We developed personalized head models from the structural MRI scans of 57 older adults (age ≥ 65 years) who participated in one of two previous tDCS trials (NCT03814304 and NCT04295798). Both studies enrolled healthy men and women aged 65 years or older, without overt neurological or psychiatric illness. All participants provided written informed consent as approved by the Hebrew SeniorLife Institutional Review Board. T1w (GE scanner, 3D SPGR sequence, acquired with 32 channel head coil) and T2w (3D CUBE sequence) MRIs were segmented into the skin, skull, air cavities, cerebrospinal fluid (CSF), gray matter (GM), and white matter (WM) using SimNibs (v3.2.1) (Saturnino, Puonti, et al., 2019) (Figure 1a, first column). All segmentations were inspected and corrected manually if deemed necessary. The most common corrections were due to errors in the CSF and bone over/under-segmentation.

**Figure 1:**
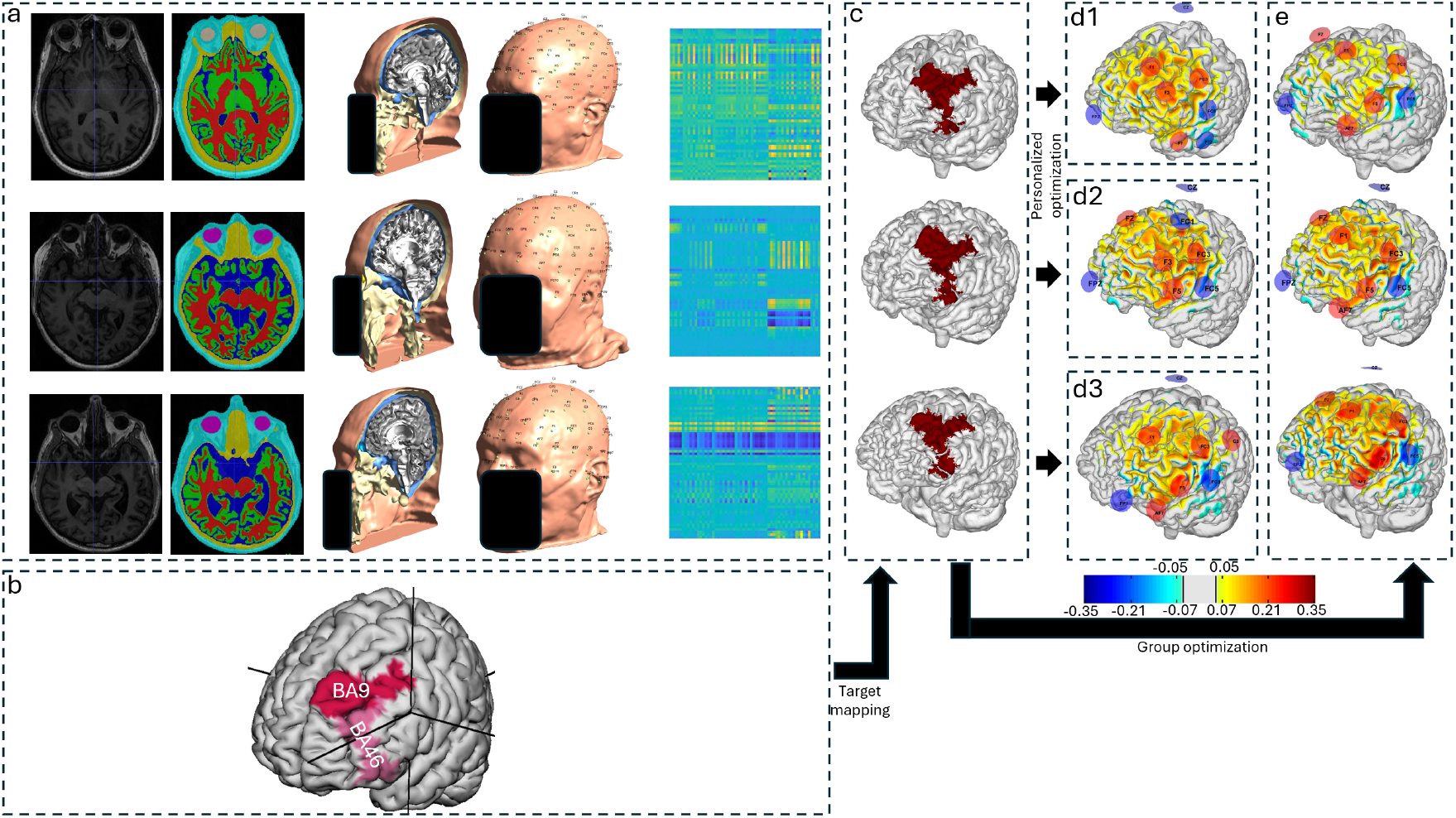
The modeling steps performed in this study. (a) From left to right: Segmentation of the structural head MRIs into the different tissues, construction of a 3D head model for finite element calculation, mapping of the 10-10 EEG system electrode positions to the surface of the scalp, and calculation of the lead-field matrix for every bipolar combination of electrodes with Cz as a cathode. (b) The left dorsolateral prefrontal cortex (lDLPFC) target is defined in the head surface of the MNI template. The target comprises Brodmann areas 9 and 46. (c) The lDLPFC target in the grey-matter (GM) surface of each individual head model after mapping from the MNI space to the participant’s native space. (d) Distribution of the En-field in the cortical surface (GM surface) of each participant, induced by subject-specific protocols targeting the lDLPFC, was obtained by performing a personalized optimization of the normalized error with respect to no intervention (NERNI) for each participant. (e) Same as d, but now for the group-optimized protocol obtained from maximizing the average of NERNI across all participants. A common color scale was used for all the plots of En (in units of V/m). Scalp reconstructions have been anonymized.

For each participant, a finite element mesh was built, and 10-10 EEG system positions were identified on the scalp surface (Figure 1a, second column). The positions were then manually inspected for errors in the registration of the electrodes, which led to the removal of two of the participants. The lead-field matrix for the E-field component perpendicular to the cortical surface (*E*_*n*_) was then calculated (Figure 1a, third column). Models of Ag/AgCl electrodes (modeled as 1 cm radius cylinder of conductive gel, with a height of 3 mm) were added to the head mesh, and tissues modeled with specific conductivities (Mercadal et al., 2022): 0.33 S/m, 0.008 S/m, 1.79 S/m, 0.40 S/m, 0.15 S/m respectively for each one of the tissues mentioned before. The gel was modeled with a conductivity of 4.0 S/m. Each column of the lead-field matrix (***K***) contains En per node (V/m) for each bipolar montage with a fixed cathode (Cz, -1 mA).

Similar methods were used to generate the head models for the templates Colin (Miranda et al., 2013) and ICBM152 (ICBM 2009c Nonlinear Asymmetric template) (V. Fonov et al., 2011; V. S. Fonov et al., 2009).

### Montage and current optimization

The *Stimweaver* algorithm (Ruffini et al., 2014) was used to maximize a fitness relative to no intervention (NERNI), defined as the least-squared difference between a weighted *E*_*n*_ induced by the montage and a weighted target 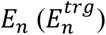,

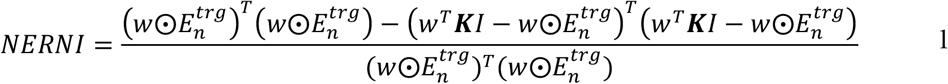

where *w* is an array of weights (*N*_*mesh*_ × 1 array, where *N*_*mesh*_ is the number of mesh nodes in the cortical GM surface), 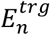 is the target *E*_*n*_-field array (dimensions of *N*_*mesh*_ × 1, in units of V/m), ***K*** is the lead field matrix (*N*_*mesh*_ × *N*_*electrodes*_ - 1, in units of V/m per mA of injected current, *N*_*electrodes*_ is the number of electrode positions defined in the scalp), and *I* is the array with electrode currents in mA (Cz excluded, as its current is defined implicitly).

NERNI is a scalar fit parameter that measures the similarity of *E*_*n*_ to 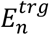. It is equal to one for a perfect fit and a large negative number when the fit is poor.

Targets were defined by mapping Brodmann areas 9 and 46 (left dorsolateral prefrontal cortex, lDLPFC) from an MNI template to the GM surface of each participant (Figure 1b). The optimization constrained the maximum current at any electrode to 2.0 mA and the total injected current to 4.0 mA using a genetic algorithm to limit the montage to eight electrodes. The target *E*_*n*_ was set to 0.25 V/m in the lDLPFC, with a weight of 10 indicating an increase in cortical excitability according to the lambda-E model for interactions of the E-field with neurons (Ruffini et al., 2014). In this model, the effects of stimulation on cortical excitability are due to the interaction of the normal component of the E-field with large pyramidal cells in the cortex. When *E*_*n*_ points to the cortical surface (positive *E*_*n*_ in the convention followed in this work), the soma of pyramidal cells is depolarized, which leads to an increase in cortical excitability. The negative *E*_*n*_ values have the opposite effect. This model appears to be representative of the effects of weak E-fields on neuronal membranes (Galan-Gadea et al., 2023). In the other areas, the target *E*_*n*_ was set to 0 V/m (no effect), with a lower weight of 2.

Optimization was conducted using a genetic algorithm, as explained in more detail by (Ruffini et al., 2014). Initially, a population representing combinations of electrode positions was created randomly. Then, for each combination (DNA string), SLSQP (sequential least squares programming optimization) was used to determine the currents that maximized the NERNI. The best members of the population were then selected, and genetic operators were used to determine the next population. This process was repeated and stopped when the improvement in the best NERNI was less than a specified threshold for five consecutive generations of the genetic algorithm. The method was implemented in Python, using DEAP and SciPy as the main libraries.

Protocol optimization was performed for the participants individually (Figure 1d 1-3). Additionally, group optimizations (Figure 1e) were conducted using a leave-one-out approach: for every participant a group optimization was conducted using the remaining 53 participants (templates excluded). The group optimization is similar to the single participant optimization, but the objective function is the arithmetic mean of NERNI for each of the participants included in the group (*N*_*sg*_),

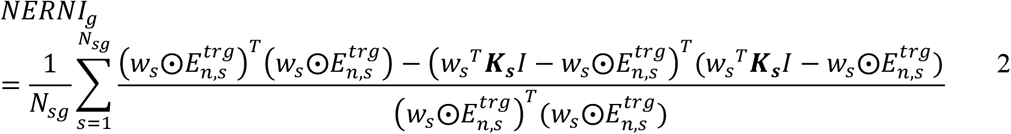

Similar methods were used to maximize *NERNI*_*g*_.

An additional metric was used to evaluate the performance of the montage, the surface average of *E*_*n*_ (< *E*_*n*_ >),

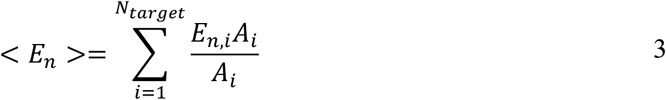

Here, *N*_*target*_ is the number of nodes in the main target region (lDLPFC) in the GM surface, *E*_*n, i*_ is the *E*_*n*_ induced at node *i*, and *A*_*i*_ is the area of the node (the sum of the areas of all triangles that share the node, divided by 3).

### Calculation of anatomical features

To test whether the anatomical features of the head models were associated with the NERNI of a protocol, we calculated several global anatomical features,

- Axial perimeter: Geodesic distance along the scalp between Nz-LPA-Iz-RPA (L/RPA: left/right pre-auricular points) in mm
- Sagittal perimeter: Geodesic distance along the scalp between Nz-Cz-Iz (mm)
- Coronal distance: Geodesic distance along the scalp between LPA-Cz-RPA (mm)
- Volumes of tissues: Volumes of the scalp, skull, CSF (excluding ventricles), GM and WM in mm^3^

Figure 2 illustrates the measurement of distances for one participant. The distance features can be assessed without the need for MRI (these distances can be obtained from the head of the participant with a measuring tape). The volume features require an MRI and at least a segmentation of the different tissues. The distances were measured in *Python* with the *Pygeodesic* library, using the triangulated scalp surface of each participant. To calculate the volumes, the *MSH* files with the finite element meshes of each participant were loaded into *MATLAB* (v2018a). The volume of each tetrahedron comprising each tissue was then calculated and summed.

**Figure 2:**
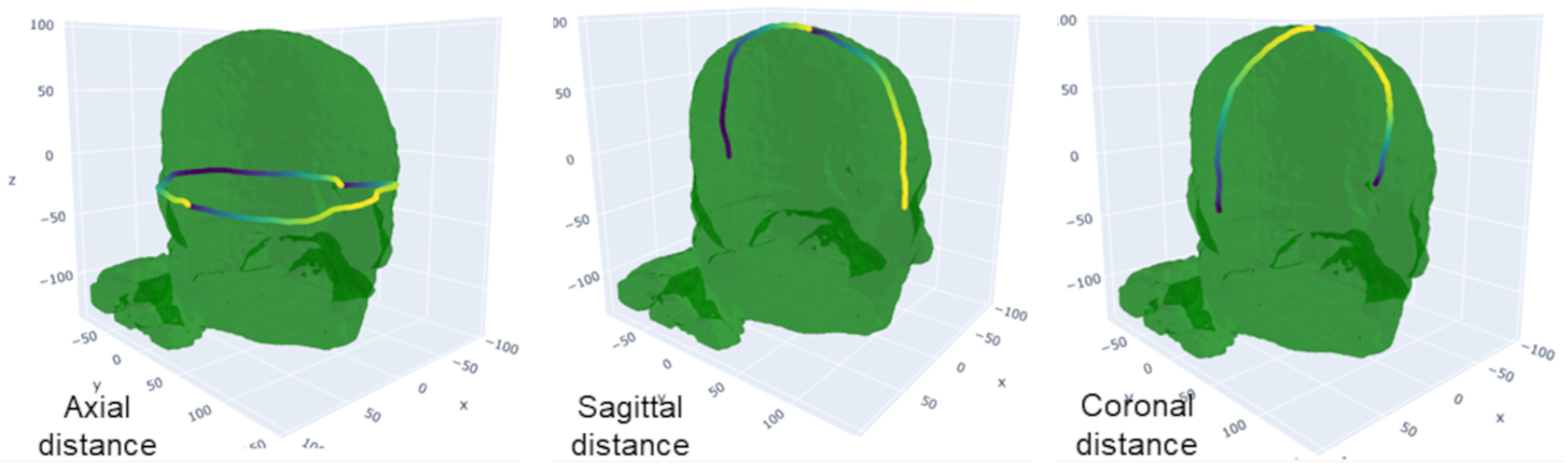
Different reference distances measured along the scalp of one of the participants. The different lines are determined by connecting reference anatomical points along the scalp of the participant: Nz (nasion), Iz (inion), R/LPA (right/left pre-auricular points).

### Statistical tests

All statistical tests were performed in *Python* (v3.7.9) with in-house functions that used *Sklearn* and *Scipy Python* libraries. Multilinear regression models were performed using with the *Statsmodel*.*api* library in *Python*.

## Results

### Personalized and group montages perform better than templates

Across all participants (see Figure 4a), the personalized protocol performed best in terms of fit (NERNI=0.21±0.03, average ± standard deviation across all participants). Both head model templates (Colin and ICBM152) performed worse than the personalized approach: 0.16±0.03 and 0.18±0.03, respectively, for Colin and ICBM152. The group approach performed better than the templates but worse than the personalized protocol: 0.19±0.03. To compare the significance of these differences, we ran a Kruskal-Wallis test, which indicated significant differences between groups (p-value<0.001). A post-hoc pairwise Dunn test with Bonferroni correction was then performed. The results are shown in Figure 4. Statistically significant differences were observed between the personalized protocols and the other cases, as well as between the group approach and the Colin template. The differences between the group protocols and ICBM152 template protocols were not statistically significant for this population size.

**Figure 3:**
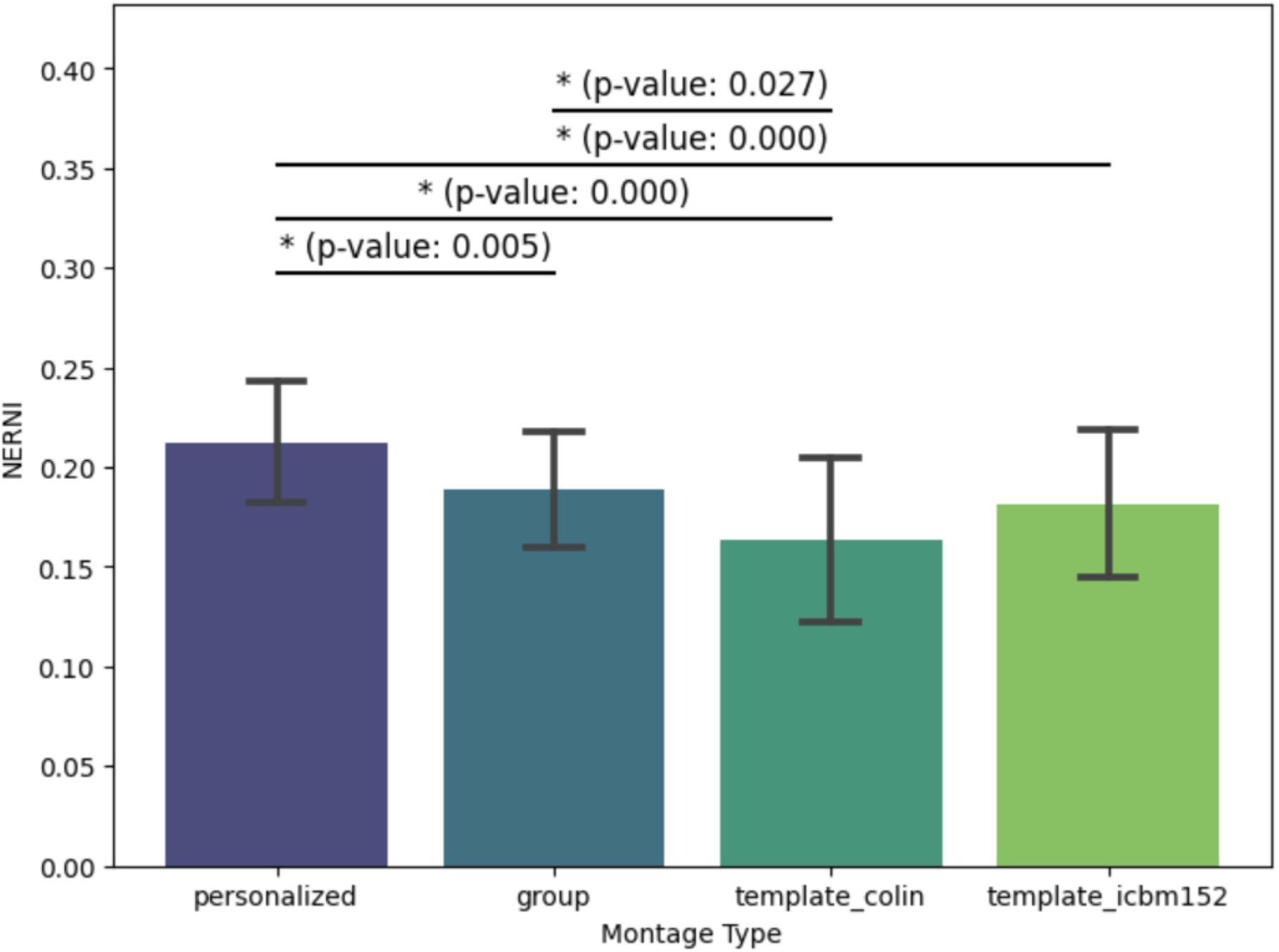
Average and standard deviation of the NERNI distributions obtained in the test participants when the protocols are obtained from different sources. The p-values were obtained with a post-hoc pairwise Dunn test with Bonferroni corrections. Only statistical significant differences (p-value<0.05) between the groups are shown.

**Figure 4:**
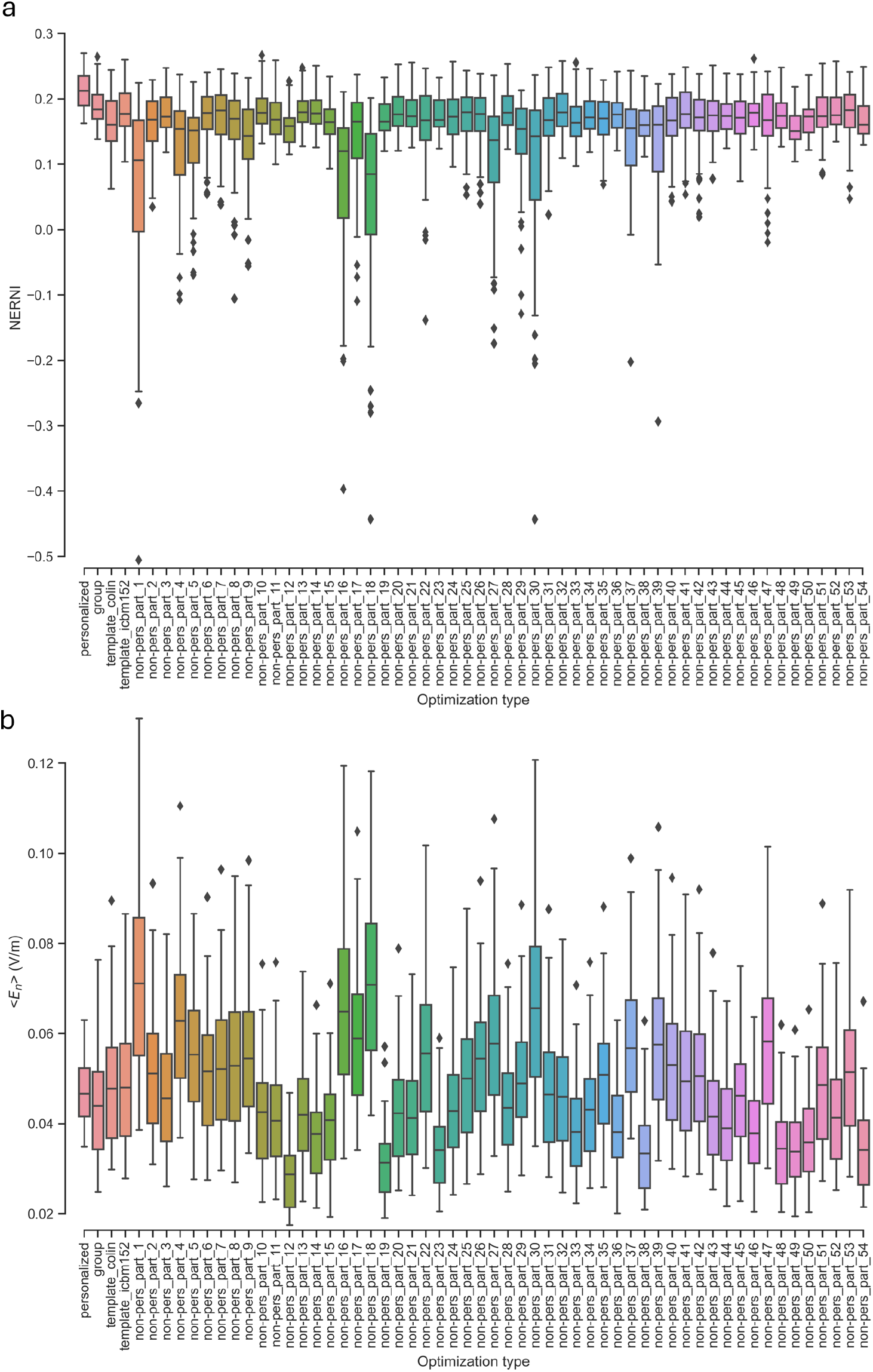
Distribution of the normalized error with respect to no intervention (NERNI) (a) and < *E*_*n*_ > (b) induced by protocols derived from different approaches: personalized, group, Colin-based template, ICBM152-based template, and templates based on each one of the biophysical head models included in the analysis (non-pers_part_<*id*>).

An example of these differences can be seen in Figure 5, for the case where the group approach performed the best (Figure 5a-d) and for the case where it performed the worst (Figure 5e-h). It can be seen that the protocols that result in the lower NERNI have a less focal En-field distribution, resulting in a worse fit to the target map.

**Figure 5:**
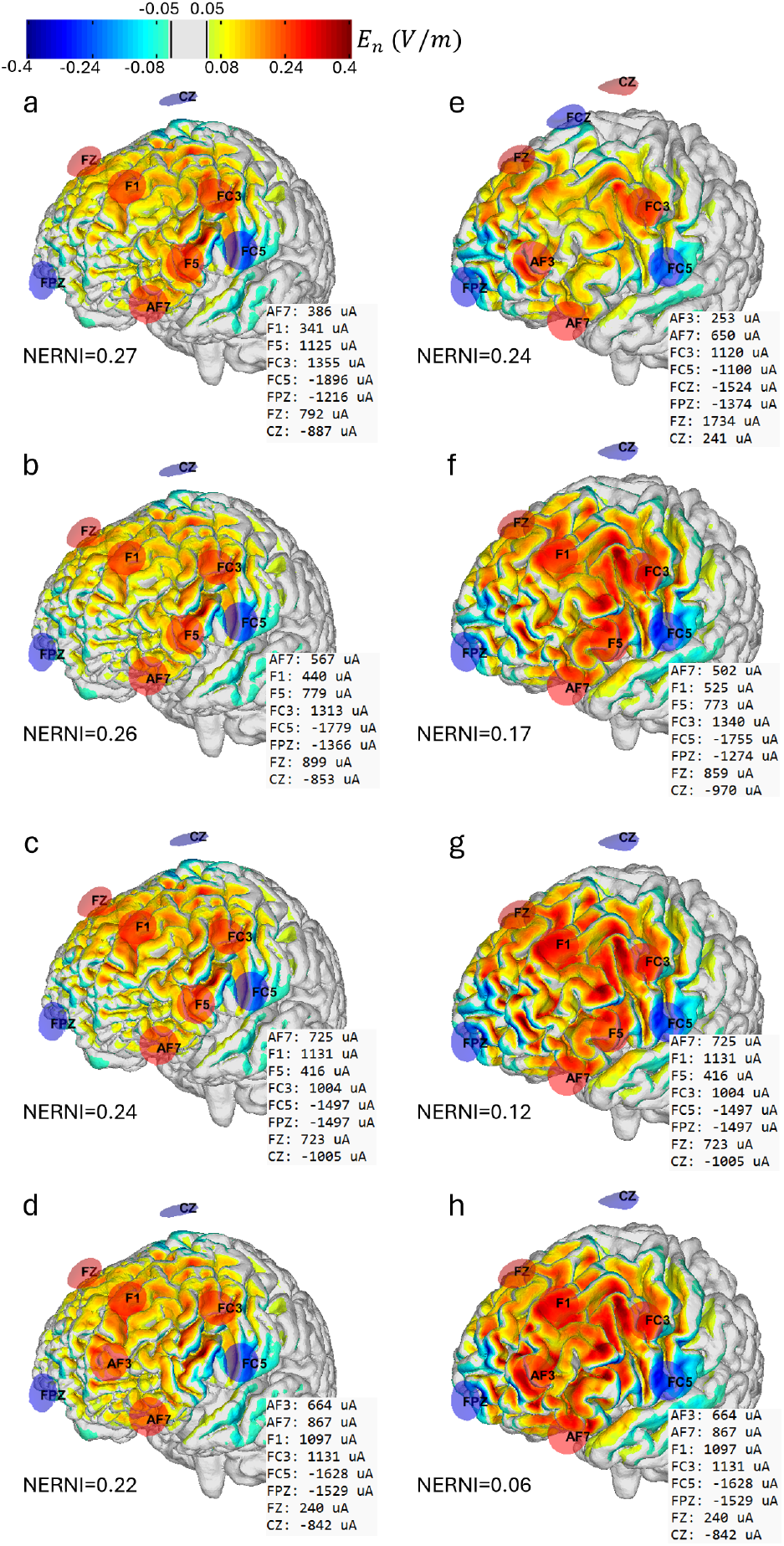
Distribution of the normal component of the E-field in the cortical surface of the participant where the group approach performed the best regarding the normalized error with respect to no intervention (left column) and for the participant where it performed the worst (right column). The figures show, in a common color scale (in units of V/m), from top to bottom: the personalized protocol, the group-optimized protocol (leave one out approach), the protocol obtained with the ICBM152 template, and the protocol obtained with the Colin template.

### Using arbitrary individual models vs standard templates

Using a non-personalized template derived from the head model of one participant led to variable results, as shown in Figure 4a. When compared to other approaches, these non-personalized templates always performed worse than the personalized templates (p-value < 0.05). The group-optimized approach always resulted in an average NERNI higher than those obtained with these non-personalized templates, but the results were only significant for 38 head models. Both the Colin and the ICBM152 template performed worse than the group montage: the average NERNI indicated that the ICBM152 template was only better than the non-personalized templates in 48 cases (24 of which were significantly different), and the Colin template was only better than the non-personalized templates in 20 cases (9 of which were significantly different).

When examining the distribution of the < *E*_*n*_ > in the target region, we can see that there are considerable fluctuations in the average value, depending on the approach used to generate the protocol (Figure 4b). A larger < *E*_*n*_ > on target did not predict good performance for the NERNI. For instance, the montages from participants 1, 16, and 18 had a large average < *E*_*n*_ > when evaluated in the other participants (albeit with larger variability as well), but the average NERNI was among the lowest. The relationship between NERNI and < *E*_*n*_ >, shown in Figure 6, shows that the entire data combination is well fitted by a quadratic regression model (R^2^=0.46, with a p-value of fit coefficients of 0.567 for the intercept, non-significant, but highly significant for the first and second-order terms: p-value < 0.0001). This indicates that very low and very high < *E*_*n*_ > values result in poor NERNI. The personalized approach restrains the protocols in a region where there is a linear increase in NERNI with < *E*_*n*_ > (Figure 6a). The group approach, ICBM152, and Colin template-derived protocols performed increasingly worse, as shown by the larger spread in < *E*_*n*_ > and NERNI values (Figure 6a, b, and c). The non-personalized templates derived from the head models of the participants resulted in a large spread of < *E*_*n*_ > and NERNI (Figure 6d).

**Figure 6:**
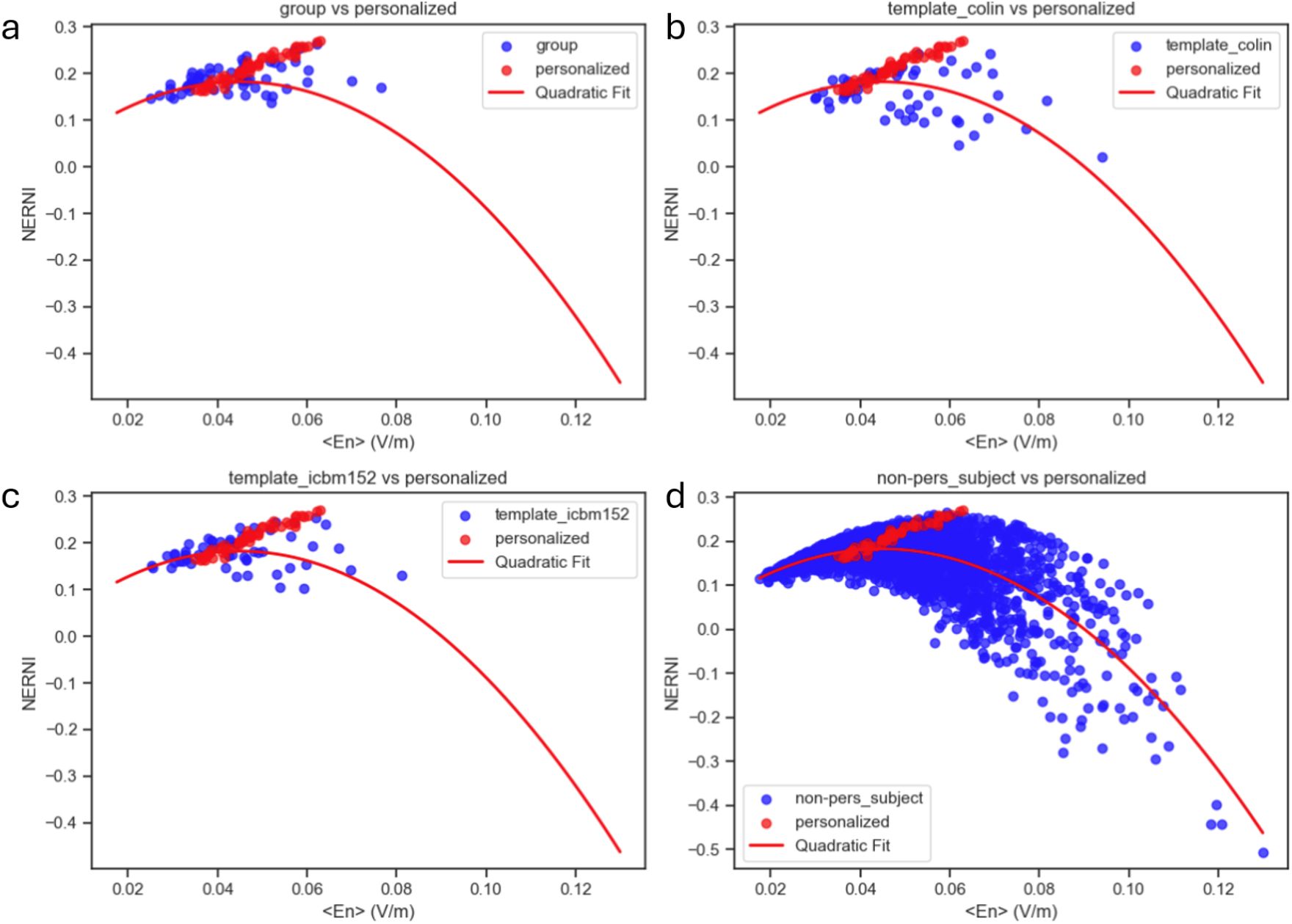
Relationship between the normalized error with respect to no intervention (NERNI) and < *E*_*n*_ > on the. left dorsolateral prefrontal cortex. The solid line indicates the fit to the data, and the red points indicate the data points obtained from the personalized approaches. The blue points indicate the other different optimization approaches: group (a), Colin-based template (b), ICBM152-based template (c), and templates based on each one of the biophysical head models included in the analysis (d).

### Anatomical features drive variance in template average En and NERNI

For the data derived using non-personalized templates obtained from the head models of the participants, we investigated the correlation of differences in anatomical features between the template head model and those of the participants in which the template was evaluated. Linear regressions of < *E*_*n*_ > against each tested individual feature showed that some of them significantly explained a small amount of the variability in NERNI (p-value < 0.05). The features with the highest correlation (see Figure 7) with < *E*_*n*_ > were the differences in the perimeter along the sagittal (*R*^2^ = 0.16) and coronal planes (*R*^2^ = 0.20), the difference in scalp volume (*R*^2^ = 0.14), and the difference in white matter and gray matter volume, normalized by the sum of the volumes of all tissues (*R*^2^ = 0.11 and *R*^2^ = 0.21).

**Figure 7:**
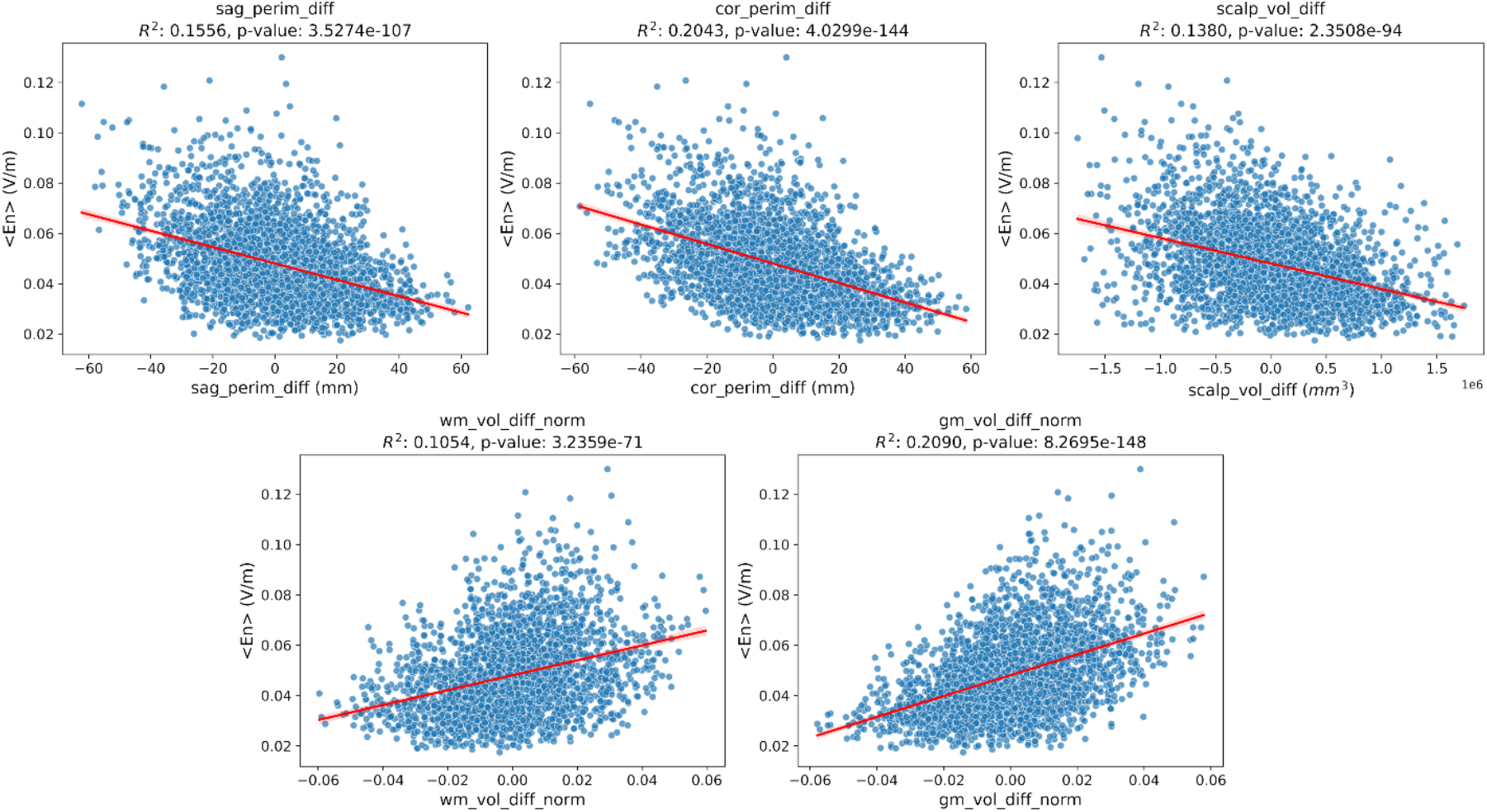
Linear regression of < *E*_*n*_ > vs the difference in anatomical features between the template used to evaluate the protocol (non-personalized template, based on the biophysical head model of one of the participants) and the participant’s anatomical features. Features from left to right: difference between perimeter of the heads along an sagittal/coronal plane (sag/cor_perim_diff), difference between the volumes of the scalp (scalp_vol_diff), and difference between the volumes of the WM/GM normalized by dividing them by the sum of the volumes of all the tissues (WM/GM _vol_diff_norm).

To perform the same analysis to NERNI, and since the previous results showed that NERNI could be fit to < *E*_*n*_ > values using a 2^nd^ order polynomial, we used a linear regression model taking as features all second-order terms in every possible combination of two features (feature-1, feature-2, feature-1×feature-2, feature-1^2^ and feature-2^2^) resulting in the plots shown in Figure 8 (only correlations with R^2^ larger than 0.1 are shown). The interactions that explain most of the variability in the data typically involve a perimeter feature (the differences in axial/sagittal/coronal distance, normalized by the sum of all three distances) and a volume feature (the difference in the scalp, skull, GM, WM, normalized, or not, by the sum of all volumes).

**Figure 8:**
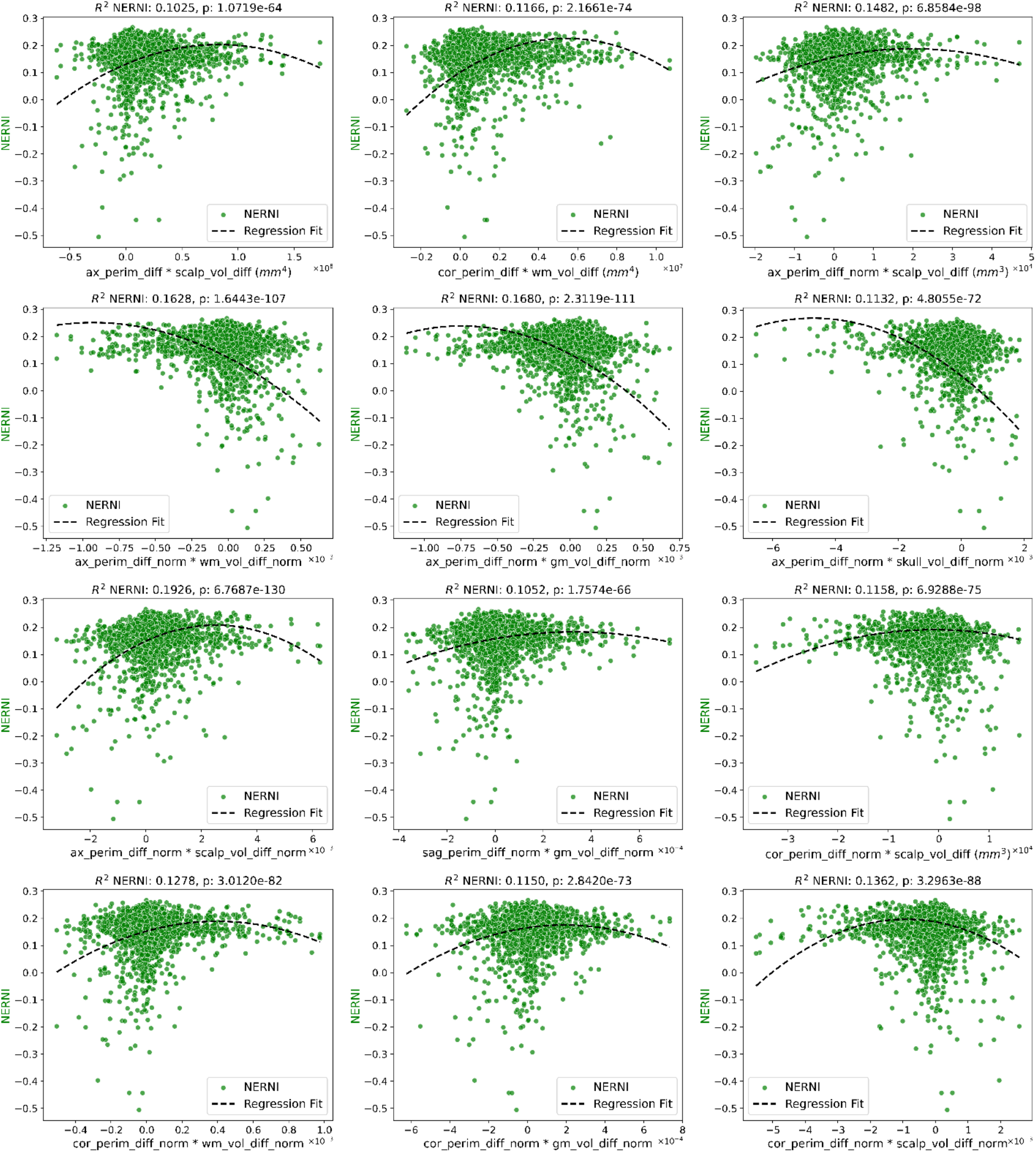
Linear regression of NERNI using as features all linear and second order terms resulting from combining differences in anatomical features between the template used to evaluate the protocol (non-personalized template, based on the biophysical head model of one of the participants) and the participant’s anatomical features. Only interactions resulting in *R*^2^ values larger than 0.1 are shown. The relevant features include: ax/cor_perim_diff (scalp’s perimeter measured along the axial and coronal planes), ax/cor/sag_perim_diff (scalp’s perimeter measured along the axial, coronal and sagittal planes, and normalized to the sum of all 3 perimeters), scalp/wm_vol_diff (volume of the scalp/WM volume), and scalp/wm/gm/skull_vol_diff_norm (volume of the scalp/WM/GM/skull, normalized by the sum of the volumes of all the tissues).

Since several of these features correlated with < *E*_*n*_ >/NERNI, we explored multilinear regression models to assess their predictive power. To evaluate the practical applicability of these models in a realistic scenario, we performed a leave-one-subject-out approach: for each subject, we removed all data from that individual, fit the multilinear regression model using the remaining participants, and then predicted the average < *E*_*n*_ >/NERNI induced by the subject’s personalized protocol in the other participants. Importantly, these predictions are based only on anatomical features.

This approach resulted in a strong correlation with the observed < *E*_*n*_ > values (*R*^2^ = 0.52, p-value=9.6×10^-10^), as shown in Figure 9a. While some of the anatomical features included in the previous analysis require MRI-based segmentation, others, such as the perimeters along the axial, sagittal, and coronal planes, can be measured without MRI access. When restricting the regression model to only these perimeter-derived features, we still found a statistically significant correlation with the observed < *E*_*n*_ > values (*R*^2^= 0.25, p-value=1.1×10^-4^; Figure 9b).

**Figure 9:**
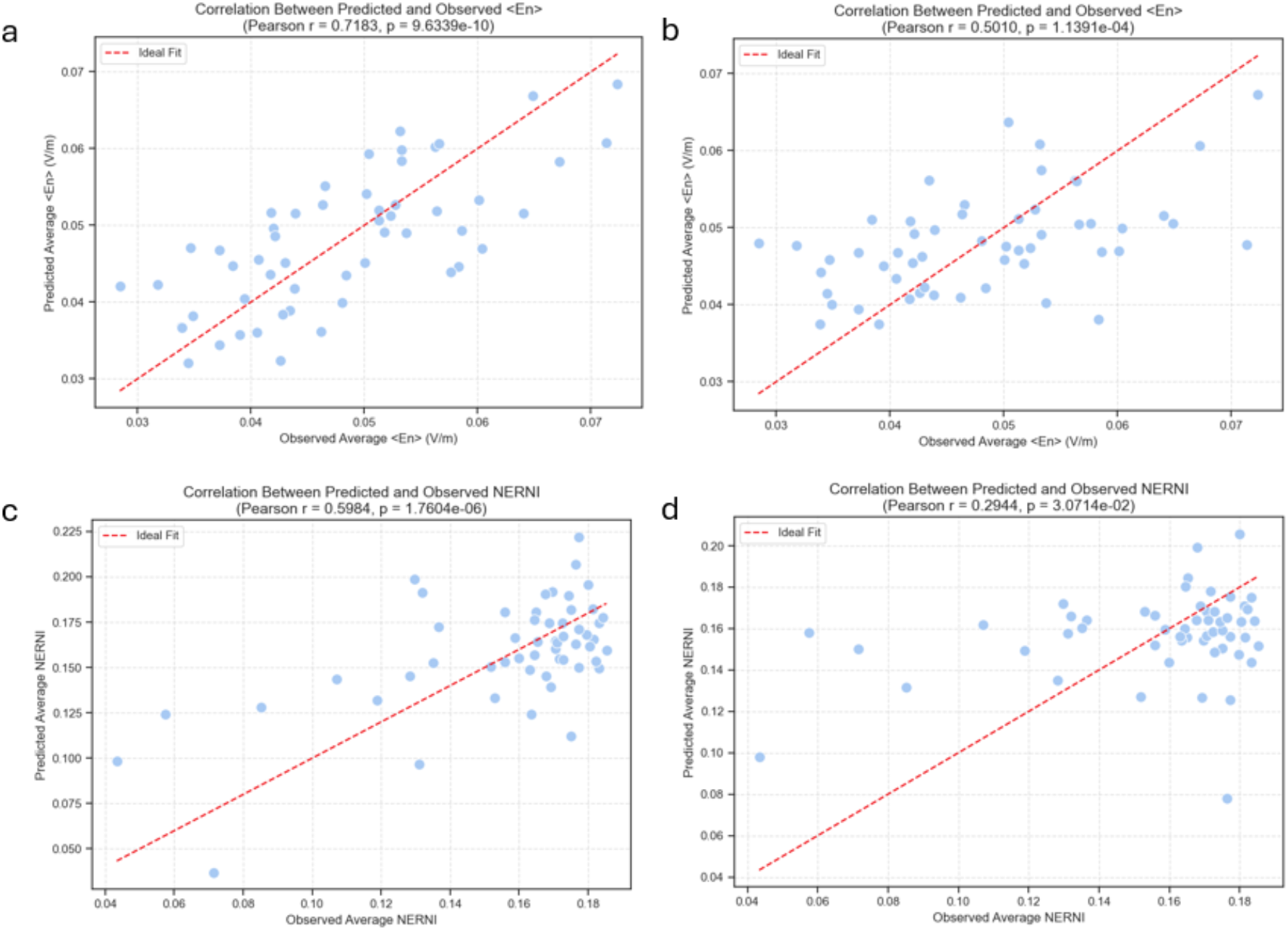
Plot of predicted and observed average < *E*_*n*_ > and NERNI (top/bottom row, respectively) for every template, derived from the (other) head models of the participants. (a/c) Model fit with all anatomical features (perimeters and volumes); (b/d) Model fit with only the perimeters of the scalp.

Extending the model to include second-order combinations of anatomical features was also successful in predicting NERNI values for both the model with the tissue volume features (Figure 9c, *R*^2^ = 0.36, p-value=1.8×10^-6^) and that with only the perimeter-derived features (Figure 9d, *R*^2^ = 0.09, p-value=3.1×10^-2^). Because there is a high collinearity between some of these features, we also tried an approach where PCA was performed before the fit, and then the fit was applied to a selection of the principal components. This resulted in slightly better results: *R*^2^ = 0.43, p-value=6.4×10^-8^ (model with all the features, and keeping 97 principle components for the fit); *R*^2^ = 0.09, p-value=2.6×10^-2^ (model with only perimeters, and keeping 17 components).

## Discussion

Our results indicate that protocol optimization algorithms can benefit from group-based approaches instead of relying on a single head model in cases where personalized head models cannot be obtained. On average, the group approach produced NERNI scores that were consistently higher than those obtained when using a particular head template, either standard templates (such as those derived from Colin or ICBM152) or templates obtained from the same population used in the study. The differences were not always statistically significant, which can be attributed to our limited sample size. The biggest advantage of our method is that it removes the need to select a “good” common template, which is complex, especially in populations with a broad range of demographic characteristics.

The correlation observed between the NERNI scores and some anatomical characteristics (i.e., the volume of the head tissues and the perimeters of the head along the reference coordinate planes) indicates that there is room to improve the results of group optimization approaches by selecting subjects in an available data pool with anatomical characteristics that predict the highest NERNI scores in the target population. Our results showed that models can be built to make these predictions based on the tested anatomical characteristics of the participants’ heads. Models that use only anatomical features that can be obtained without MRI are particularly interesting. This model performs worse than the more complex model, but our results still show some predictive power. Future work should test whether choosing subjects for the group used in the optimization based on these model predictions can improve the results. Another potential avenue for exploration lies in extending the anatomical characteristics considered. Of particular relevance, given previous work, is the local tissue thickness (Mosayebi-Samani et al., 2021; Opitz et al., 2015), which was not explored here but may further increase the predictive power of these models.

This study focused on NERNI, a more precise and complex metric than the more commonly adopted dose metrics, such as the average E-field on the target. In particular, given the relationship between these two metrics, a very high average En-field on the target (< *E*_*n*_ >) does not necessarily result in a high NERNI, especially if it comes at the cost of less focality, that is, a high En-field in regions of the target map that were set to a low target En. The balance between focality and E-field intensity on the target is encoded in the selected weights and target *E*_*n*_ value used in the optimization: a higher target *E*_*n*_ and a higher ratio of weights on the target to off-target results in montages with higher < *E*_*n*_ >, but also with worse focality.

The correlation between NERNI and stimulation effects has not yet been quantified, but several studies have reported positive effects of NERNI/ERNI-based optimizations: (Daoud et al., 2022; Fischer et al., 2017; Kaye et al., 2021; Ruffini et al., 2024; Sprugnoli et al., 2021; Zhou et al., 2021). Furthermore, a linear relationship has been shown between performance on some tasks, such as dual-task error (Manor et al., 2018) and < *E*_*n*_ > following stimulation of the lDLPFC (Salvador et al., 2022). Given the linear relationship observed between NERNI and < *E*_*n*_ > in the optimized montages (see Figure 6a), it is likely that NERNI also correlates with stimulation effects. However, the effectiveness of NERNI, hinges on the relevance of the target map for the specific application. It should be noted that group-based optimization can be performed regardless of the objective function of the optimization, and the methods reported in this paper are still valid.

This group-based approach was employed successfully in a trial on depression targeting the lDLPFC (Ruffini et al., 2024), using a cohort of patients selected to have an age range matching that of the target population. Another variation of this approach is currently being tested, in which a group optimization based on a cohort of patients with similar demographic characteristics is first performed to obtain a common pool of electrodes for every subject, and then the currents are personalized either fully (NERNI minimization, using the lDLPFC as a target, as is currently being performed in the trial: NCT06821568 ), or by scaling them via correlations obtained from the head perimeters (more details in Mencarelli et al, 2025, in prep, where the targets were selected due to their relevance in Alzheimer’s onset and progression). These applications highlight the potential of group-based optimization in real-world settings and emphasize the need for further refinement in selecting the most representative head models.

The fact that this study focused only on one target area should not be perceived as a limitation, as the general methodology is applicable to any target and, qualitatively, the results should not change. The hyper parameters used for the optimization (i.e., the weights and target En values) potentially have a stronger influence on the results. In particular, we observed that with a target En of 0.25 V/m, the participant specific optimized protocols tended to use all available total injected currents, for the majority of cases. This would not be the case for lower target En values (for instance, 0.10 V/m). This would introduce a larger gap in NERNI between the optimized protocols and the group and/or template-based approaches. In this scenario, the advantages of the group over template-based approaches are likely to be higher. The biggest advantage of lowering the target En would be to obtain a better focality of stimulation (i.e., restricting the montage to the target area) at the cost of average En on the target.

The results of the current study were based on a relatively small sample size. Future studies should focus on leveraging large databases with head structural MRIs suitable for biophysical modeling. This would increase the predictive power of the observed correlations and enable the development of strategies for stratification according to relevant anatomical/demographic features, which could mitigate the increased inter-subject variability typically associated with larger cohorts.. Additionally, this study utilized standard conductivities for the different segmented tissues. As more data emerge showing considerable inter-subject variability in tissue conductivity and that such variability induces meaningful effects on E-field distribution (Antonakakis et al., 2020; Saturnino, Thielscher, et al., 2019), it is important to take this into account in the optimization strategies. As more information is available about the expected distribution of tissue conductivities across the population, this information can be considered by expanding the database with the same biophysical head models with different conductivity values for the different tissues (see Figure 10).

**Figure 10:**
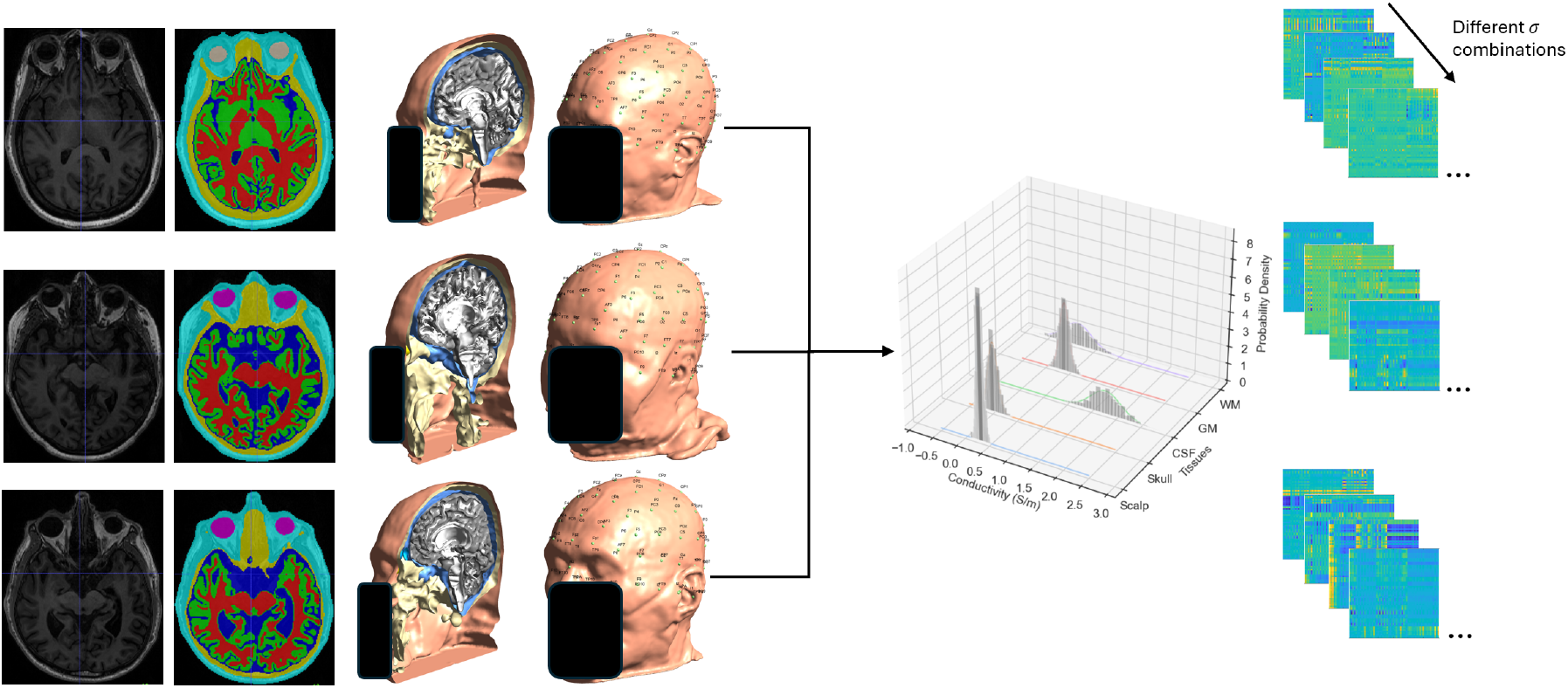
Extension of the group optimization to include repetitions of subjects with different tissue conductivity combinations based on known *a priori* distributions.

## Acknowledgements

This work was supported by European Unions Horizon 2020 research and innovation programme under grant agreement No. 101017716 (Neurotwin), by the European Research Council (ERC Synergy Galvani) under the European Union’s Horizon 2020 research and innovation programme Grant Number 855109, and by funding from the National Institutes of Health (R01AG059089; 1R21AG064575; 5K01AG075180).

## Conflicts of interest

Dr. Ricardo Salvador works for Neuroelectrics Barcelona SLU, a company developing computationally-driven brain stimulation solutions. Dr. Giulio Ruffini works for and is a co-founder of Neuroelectrics and holds several patents in model-driven non-invasive brain stimulation.

## Notes

### Competing Interest Statement

GR and RS work for Neuroelectrics Barcelona SLU, a company developing computationally-driven brain stimulation solutions. GR is a co-founder of Neuroelectrics.

